# Tilt In Place Microscopy (TIPM): a simple, low-cost solution to image neural responses to body rotations

**DOI:** 10.1101/2022.09.11.507428

**Authors:** Kyla R. Hamling, Yunlu Zhu, Franziska Auer, David Schoppik

## Abstract

Animals use information about gravity and other destabilizing forces to balance and navigate through their environment. Measuring how brains respond to these forces requires considerable technical knowledge and/or financial resources. We present a simple alternative: Tilt In Place Microscopy (TIPM). TIPM is a low-cost and non-invasive way to measure neural activity following rapid changes in body orientation. Here we used TIPM to study vestibulospinal neurons in larval zebrafish during and immediately after roll tilts. Vestibulospinal neurons responded with reliable increases in activity that varied as a function of ipsilateral tilt amplitude. TIPM differentiated tonic (i.e. sustained tilt) from phasic responses, revealing coarse topography of stimulus sensitivity in the lateral vestibular nucleus. Neuronal variability across repeated sessions was minor relative to trial-to-trial variability, allowing us to use TIPM for longitudinal studies of the same neurons across two developmental timepoints. There, we observed global increases in response strength, and systematic changes in the neural representation of stimulus direction. Our data extend classical characterization of the body tilt representation by vestibulospinal neurons and establish TIPM’s utility to study the neural basis of balance, especially in developing animals.

**Significance Statement:** Vestibular sensation influences everything from navigation to interoception. Here we detail a straight-forward, validated and nearly-universal approach to image how the nervous system senses and responds to body tilts. We use our new method to replicate and expand upon past findings of tilt sensing by a conserved population of spinal-projecting vestibular neurons. The simplicity and broad compatibility of our approach will democratize the study of the brain’s response to destabilization, particularly across development.

## Introduction

Animals must transform forces acting on the head/body into commands to stabilize gaze/posture, orient, navigate, and regulate physiology (Angelaki and Laurens, 2020; Chen et al., 2021; Daltorio and Fox, 2018; Yates et al., 2013; Yoder and Taube, 2014). Both sensory organs and bodies change as animals develop and age. Studying neuronal representations of forces – particularly over time – presents a profound challenge: to measure brain activity while applying destabilizing forces. For over a century novel apparatus have met this challenge, primarily in studies of the vertebrate vestibular system: from observational chambers (Lowenstein and Roberts, 1949; Mach, 1886; Straka et al., 2021), translating sleds (Fleisch, 1922), and later to electrophysiology-compatible rotating swings (Adrian, 1943), modern platforms with 6 degrees of freedom (Angelaki et al., 1999; Branoner and Straka, 2018) and wireless recording (Carriot et al., 2017). Forces are usually delivered sinusoidally, conferring mechanical advantages and facilitating linear systems analysis and modelling of neuronal/behavioral responses (Laurens et al., 2017). While studies of vestibular processing have advanced nearly every area of systems neuroscience (Goldberg et al., 2012a), each apparatus and stimulus paradigm represents a set of necessary trade-offs. The specialized hardware used in most experiments is both a cost and knowledge barrier. Further, few existing approaches permit repeated measures of activity from the same neurons over days, key to understanding changes that accompany development (Beraneck et al., 2014; Peusner, 2014) and learning (Goldberg et al., 2012b). Here we detail and validate an approach for longitudinal measures of vestibular sensitivity.

Imaging neuronal activity by measuring changes in fluorescence of genetically-encoded calcium indicators has transformed neuroscience. The vestibular field has been comparatively slow to adopt this technology largely because imaging neurons while animals are challenged with vestibular stimuli is technically demanding. Recent efforts include microscopes that rotate (Migault et al., 2018), image a rotating sample (Hennestad et al., 2021; Tanimoto et al., 2022), and optically trap and move part of the vestibular apparatus (Favre-Bulle et al., 2018). Taken together, these studies clearly illustrate the promise of using modern microscopy and genetics to understand the vestibular system. However, each requires specialized optical and/or engineering expertise to implement, limiting their impact. Motivated by these studies, we sought to develop a complementary low-cost, straightforward means to image neural activity following body tilts.

We focused our efforts on a model vertebrate with a rich vestibular repertoire: the larval zebrafish. At four days old larval zebrafish swim freely and maintain posture and stabilize gaze (Bagnall and Schoppik, 2018). Both behaviors require central vestibular neuronal circuits (Schoppik et al., 2017), with considerable development between four and seven days (Bianco et al., 2012; Ehrlich and Schoppik, 2017, 2019). Importantly, the larval zebrafish is both transparent and genetically-accessible, facilitating measurements of calcium indicators in vestibular neurons, and/or loss-of-function assays in mutants (Whitfield et al., 1996). Finally, like all vertebrates, the larval zebrafish has a small population of vestibulospinal neurons (∼50 cells) with reliable and well-characterized responses to tilt stimulation (Hamling et al., 2021; Liu et al., 2020). Taken together, the zebrafish is a powerful model to develop new methods to investigate vestibular function.

Here we present data from a new approach to imaging neural activity in response to body rotations we call Tilt In Place Microscopy, or TIPM. TIPM allows extremely rapid (*<*6 ms) whole-body rotations toward and away from the horizon, allowing precise characterization of tilt sensitivity. We validated TIPM by characterizing the roll responses, topography, and development of larval zebrafish vestibulospinal neurons. We found that vestibulospinal neurons respond reliably to ipsilateral steps with parametrically increasing activity, consistent with prior electrophysiological measurements in fish and mammals (Peterson, 1970; Rovainen, 1979). We repeated TIPM sequentially on the same fish and found that trial-to-trial variation was likely intrinsic to vestibulospinal responses, not due to our approach/apparatus. Vestibulospinal neurons had a comparatively small response to phasic stimulation; neurons that sensed phasic components were preferentially located in the ventral lateral vestibular nucleus. The bulk of the vestibulospinal response was derived from utricular sensation. Finally we measured responses from the same neurons at two behaviorally-relevant time points, revealing increased response strength in older neurons and systematic changes in directional selectivity as neurons develop. We end with a discussion of the advantages of our method (low cost, broad compatibility, extensibility) and its limitations. Our method will facilitate investigation of neuronal responses to tilt stimulation, particularly in small model animals such as *Drosophila, Caenorhabditis, Danio, Danionella*, and *Xenopus*.

## Materials and Methods

### Fish Care

All procedures involving zebrafish larvae (*Danio rerio*) were approved by the Institutional Animal Care and Use Committee of New York University Grossman School of Medicine. Fertilized eggs were collected and maintained at 28.5°C on a standard 14/10 hour light/dark cycle. All experiments were performed on larvae between 4 and 7 dpf. During this time, zebrafish larvae have not yet differentiated their sex into male/female. Before 5 dpf, larvae were maintained at densities of 20-50 larvae per petri dish of 10 cm diameter, filled with 25-40 mL E3 with 0.5 ppm methylene blue. After 5 dpf, larvae were maintained at densities under 20 larvae per petri dish in E3 and were fed cultured rotifers (Reed Mariculture) daily.

### Fish Lines

All experiments were done on the *mitfa* -/- background to remove pigment for imaging. All larvae were labeled with *Tg(nefma:GAL4;UAS:GCaMP6s)* (Liu et al., 2020; Thiele et al., 2014). For experiments testing utricular origin of responses, *Tg(nefma:GAL4;UAS:GCaMP6s)* fish with a homozygous recessive loss-of-function mutation of the inner ear-restricted gene, *otogelin* (otog-/-), also called *rock solo*^AN66^ (Whitfield et al., 1996) were visually identified by a bilateral lack of utricular otoliths.

### Imaging Set-Up and Apparatus

Larval fish (4-7 dpf) were mounted in a small drop of 2% low melting point agarose (ThermoFisher Scientific 16520) in the center of the uncoated side of a mirror galvanometer (Thorlabs GVS0111). E3 was placed over the agarose and the galvanometer mirror was placed under the microscope.

For routine imaging, a baseline voltage was applied to the galvanometer to set it at one end of its range, allowing for up to 40° of rotation away from the horizontal plane in one direction. Stimuli were capped at ±30° to allow the experimenter to apply an additional small offset voltage to correct for slight deviations from horizontal incurred while mounting the fish, and because of steric limitations relative to the objective. All trials (3 trials per step size, 2 stimuli repeats per trial) in one direction were run with that baseline voltage manually set at horizontal, then a new baseline voltage was applied and the galvanometer was re-centered at horizontal to continue performing trials in the opposite tilt direction. Experiments with *rock solo* fish were performed with a different stimulus paradigm; in these experiments no baseline voltage was applied to the galvanometer before it was positioned at horizontal, and the maximum voltage drive was then applied during the stimulus to rotate the sample to ±20°.

For experiments where responses at horizontal were compared to responses measured directly at the eccentric angle, fish were mounted and first imaged upon return to horizontal. Then, the microscope was manually focused to allow the fish to be in focus at an eccentric angle for subsequent trials. Fish were then anesthetized by applying 0.2 mg/mL ethyl-3-aminobenzoic acid ethyl ester (MESAB, Sigma-Aldrich E10521) to the fish mounted in agarose. The fish was allowed to sit with the anesthetic for 10 minutes before imaging recommenced to ensure it had reached full effect; MESAB remained on the fish for the rest of the imaging session to keep the fish properly anesthetized. The imaging was then repeated in the anesthetized fish with the fish in focus upon return to horizontal and subsequently in focus at the eccentric angle.

Galvanometer control was done in Matlab 2019b (Mathworks, MA) using the Data Acquisition Toolbox to interface with a data acquisition card (PCIe-6363, National Instruments, TX). Custom code was written to simultaneously (1) deliver an analog waveform to control the galvanometer (2) measure the analog voltage that corresponded to the galvanometer position and (3) deliver synchronizing digital pulses to begin imaging. For all step and impulse stimuli, the galvanometer was allowed to step to/away the eccentric angle at the maximum angular velocity/acceleration achievable (Table 1). Impulse stimuli were delivered in both directions. A microscope (Thorlabs Bergamo) was used to measure fluorescence elicited by multiphoton excitation (920 nm) from a pulsed infrared laser (MaiTai HP, Newport, CA). Fast volumetric scanning was achieved using a piezo actuator (ThorLabs PFM450E) to move the objective. Each frame of the volume (416 × 64 pixels) was collected with a 1 μs pixel dwell time (18.6 frames/second). Volumes ranged from 6-9 planes in depth (6 μm between planes) to cover the entire vestibulospinal nucleus, resulting in a mean effective volume rate of 2.2 volumes per second (range 1.9-2.7 volumes/second). Fish that were imaged multiple times were imaged using the same scan settings across both sessions. The *rock solo* mutant fish and their wild-type siblings were imaged prior to the addition of the piezo actuator; in the place of volumetric imaging, for these experiments single z-planes were imaged separately (2-3 planes per fish) at a frame rate of 6.6 frames/second.

**Table 1:** Stimulus properties

|  | Unit | 10° | 20° | 30° | 10° Impulse | 15° Impulse | 30° Impulse |
| --- | --- | --- | --- | --- | --- | --- | --- |
| Maximum Angular Velocity | deg/ms | 5.3 | 6.5 | 7.5 | 5.4 | 6.1 | 6.8 |
| Average Angular Velocity | deg/ms | 4.1 | 4.9 | 5.5 | 3.9 | 4.6 | 4.2 |
| Maximum Angular Acceleration | deg/ms<sup>2</sup> | 18.6 | 20.0 | 20.2 | 19.8 | 21.2 | 20.8 |
| Average Duration of Step | ms | 2.2 | 3.9 | 5.3 | 5.0 | 5.6 | 13.0 |
| Average Duration at Eccentric Angle | - | 15 s | 15 s | 15 s | 0.3 ms | 0.4 ms | 2.5 ms |

### Data Analysis & Statistics

#### Calcium response extraction and analysis

Data analysis was performed using custom code in Matlab 2017b (Mathworks, MA). We pre-processed the imaging data with code adapted from the CalmAn package (Giovannucci et al., 2019), and then performed NoRMCorre rigid motion correction (Pnevmatikakis and Giovannucci, 2017) on our GCaMP6s signal across all concatenated trials of the same stimulus type for each fish. We then hand-drew polygon ROIs in FIJI (Schindelin et al., 2012) around each vestibulospinal cell on the maximum intensity projection of all the motion corrected frames. We imported ROIs into Matlab using Read-ImageJROI (Muir and Kampa, 2015) and extracted the mean pixel value across each ROI for all time-points of each trial, and performed normalization of the raw fluorescence trace.

For experiments imaged at horizontal, we quantified the peak calcium response to each stimulus as the mean ΔF/F in the first second after the sample returned to horizontal. For experiments imaged at the eccentric angle, responses calculated for comparison to those at horizontal were the mean ΔF/F in the last second before the sample returned to horizontal. A cell was determined to have a significant response to a stimulus if the peak calcium responses across all trials were significantly higher (one-tailed t-test, p<0.05) than the mean calcium responses during the first second of the baseline period of that same cell. For each cell, a directionality index was calculated by taking the difference between the peak calcium response to ipsilateral and contralateral 30° steps, normalized by their sum.

To calculate the sensitivity of peak calcium responses to step magnitude, we fit a line with two free parameters to the peak calcium responses from all trials of all step magnitudes in a single direction (ipsilateral or contralateral). During analyses of longitudinal calcium imaging, cells with a significant sensitivity increase/decrease between time points was defined as follows: To determine a cutoff for significantly increasing/decreasing slopes, for each fish we shuffled the peak calcium responses across all trials from both ages in the same direction, and used the shuffled responses to calculate a best-fit line with a slope. We then took the difference of the shuffled slopes between 4 and 7 dpf for each cell for the ipsilateral and contralateral direction. The cutoff for significant sensitivity change was defined as the mean of the shuffled slopes ±2 SD. In longitudinal experiments, “Early Contra Responders” were defined as cells that had a contralateral slope at 4 dpf greater than the mean + 2 SD of contralateral 4 dpf shuffled slopes.

To determine any field-of-view shifts after an eccentric step, we performed a two-dimensional normalized cross-correlation between the frame prior to the stimulus and the frame after the stimulus for each plane of the volume. These analyses were performed on frames from unprocessed volumes. Field-of-view shift in the x- and y-axis was determined by finding the position of highest correlation coefficient within the resulting matrix, and corresponding that matrix with an x- and y-axis shift in pixels (reported in text after conversion to μm). Mean correlation coefficients for each fish were calculated from the center of the correlation coefficient matrix (i.e. the correlation between the two frames without any x- or y-shift).

Correlation was evaluated using Pearson’s correlation coefficient (\*ρ*). We report slope fits and 95% confidence intervals (CI). To test for interactions between groups we use either a univariate analysis of variance test (ANOVA) or multivariate (MANOVA).

#### Normalization

When comparing activity in the same neuron measured at horizontal only (our standard imaging paradigm), we normalized the fluorescence against the mean fluorescence value in the last 5 seconds of the baseline period within each trial. When comparing activity in the same neuron measured at different angles, we used the fluorescence measured in an anesthetized condition at the angle at which the imaging was done. Our rationale is as follows: the intensity of a genetically-encoded calcium indicator reflects a number of variables, necessitating normalization. The “baseline” level is usually derived during a neutral condition with respect to the stimulus, correcting for differences in expression levels and variable imaging conditions (e.g. IR light penetration or scattering of emitted photons). Further, vestibular neurons might have different basal activity when held at eccentric positions. We assert that the fluorescent intensity measured in a given neuron in an anesthetized animal will only reflect basal expression levels and variability due to imaging conditions. Consequentially, it is an easily-accessible baseline that permits us to compare responses in the same neuron when held at different angles.

#### Vestibulospinal Cell Body Position and Roll Angle Analysis

For each fish, an additional 2-photon volumetric stack was taken with scan settings optimized for a high-signal, low-speed image (2 μs scan speed, cumulative averaging across 4 frames, 1 μm between z-planes). This stack was used for localizing the vestibulospinal cell bodies in three-dimensions, defined relative to the Mauthner cell lateral dendrite. To define these XYZ positions, we first placed reference point ROIs in FIJI at the lateral-most tip of the Mauthner cell lateral dendrite in both brain hemispheres. We then dropped point ROIs on all vestibulospinal cells that were analyzed in our calcium imaging trials, placing the ROI at the center of the cell body at the z-plane where the cell was most infocus. Using FIJI’s “Measure” tool, we measured the XYZ position of each ROI in microns, and exported this data to Excel. For each vestibulospinal cell position, we subtracted off the XYZ position of the Mauthner lateral dendrite in its corresponding brain hemisphere to convert absolute position to relative position, and then imported the relative XYZ position data to Matlab for plotting.

For calculating the roll tilt angle of each fish, we used the left and right Mauthner lateral dendrite reference ROIs to find the distance between the two hemispheres in depth (z-axis). We then calculated the average mediolateral (x-axis) distance between the Mauthner lateral dendrites (171.8 μm). We took the arctangent of the z-distance and average x-distance to calculate the roll angle for each mounting.

## Results

### Rationale, apparatus, and stimulus for Tilt In Place Microscopy

We developed a simple method (TIPM) to permit imaging of neuronal activity following body tilts. The vestibular end-organs in vertebrates detect either linear accelerations (such as orientation relative to gravity) or rotational accelerations. We reasoned that the most straightforward way to assay this activity would be to image the same volume before and after such stimulation to avoid image registration challenges and to maximize compatibility with different microscope architectures. The kinetics of fluorescent indicators of neuronal activity are slow to decay (Chen et al., 2013). Consequentially, a sufficiently rapid step back to the horizon from an eccentric orientation would produce a response with two components. The first component would reflect the steady-state activity of the neurons encoding linear acceleration due to gravity (i.e. the body’s tilt) before the step. The second component would reflect the phasic response to the step itself, if any. We refer to the stimulus that elicits these combined responses as a “step.” Complementarily, a second stimulus comprised of a rapid step to an eccentric angle and back would generate a response to an impulse of rotation, devoid of any steady-state component. We refer to this stimulus as an “impulse.” Taken together, step and impulse stimuli allow for characterization of both the tonic (i.e. steady-state body tilt) and phasic (i.e. rapidly changing) components of a neuron’s response. The key to our method is therefore delivery of rapid rotations to the preparation.

Here we used a mirror galvanometer as a platform to rapidly rotate an immobilized larval zebrafish. Mirror galvanometers are the tool of choice to steer light to particular angles for their precision and rapid response. We mounted a larval zebrafish in a small drop of low melting temperature agarose directly on the uncoated side of the mirror (Figure 1A). We could rotate the platform through nearly the full range of the galvanometer both rapidly (5.3 ms for a 30° step, (Table 1), and precisely (Figure 1B-1D).

**Figure 1:**
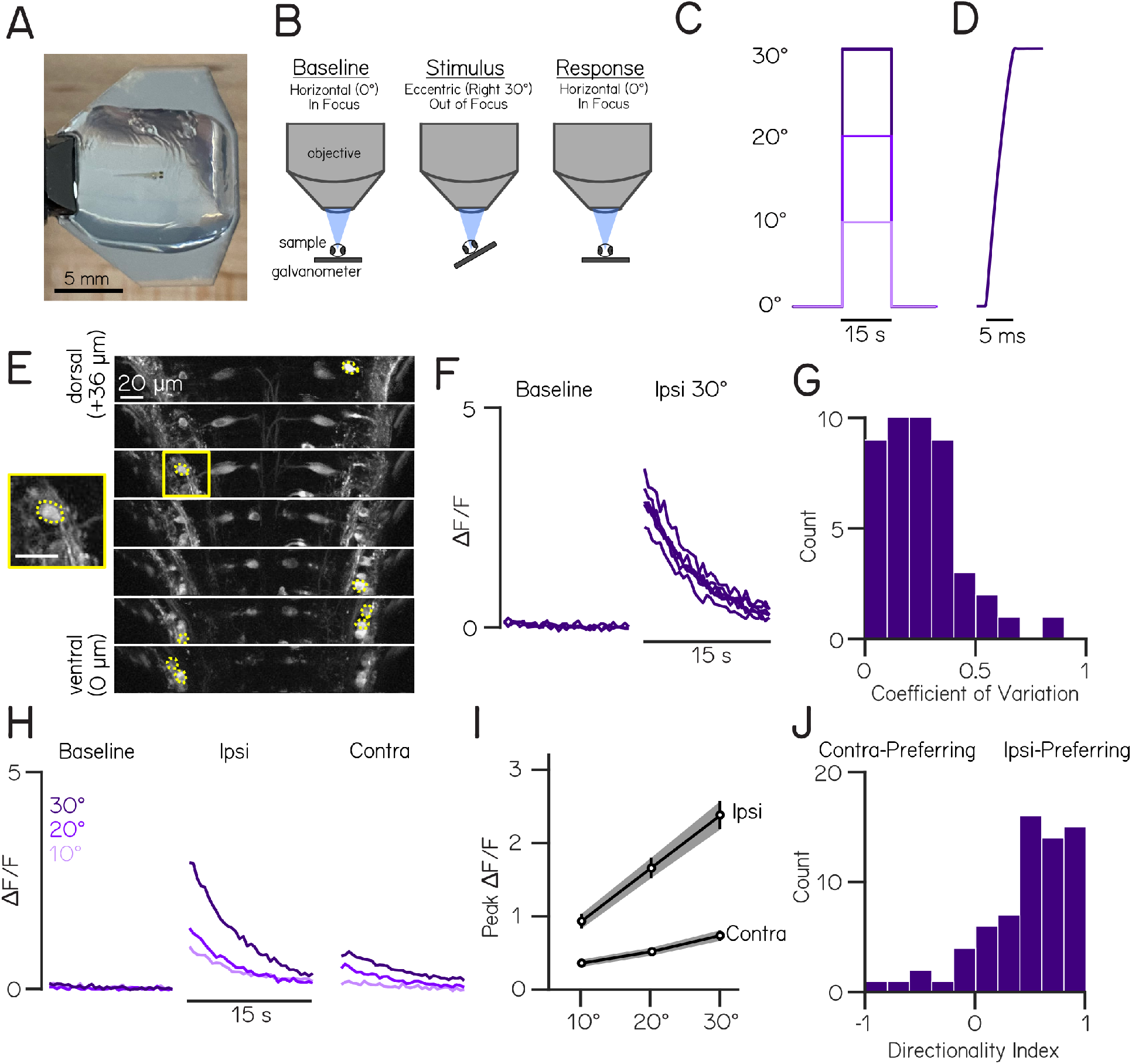
Tilt In Place Microscopy (TIPM) produces reliable, directional, and magnitude-dependent responses following roll tilts. **(A)** A 4 day post-fertilization (dpf) larval zebrafish mounted in agarose for roll stimuli on a mirror galvanometer. **(B)** Schematic of our experimental paradigm. Baseline fluorescence (used for normalization) is measured when the platform is horizontal. The galvanometer is then stepped and held at an eccentric angle (“Stimulus”) where fluorescence is not recorded, then quickly returned to horizontal whereupon fluorescent recording begins (“Response”). **(C)** Voltage-trace from galvanometer during a 10°, 20°and 30°step to the left. **(D)** Inset of feedback voltage from the galvanometer during a step to 30°.**(E)** Slices from a two-photon volume of *Tg(nefma:GAL4);Tg(UAS:GCaMP6s)* fish. Dashed yellow overlays indicate pixels that correspond to analyzed vestibulospinal neurons. Square yellow inset shows close up of a single analyzed cell (scale bar 20 μm) **(F)** Normalized fluorescence traces for all trials of one neuron during baseline and response to an ipsilateral 30° roll. **(G)** Distribution of coefficients of variation of peak ΔF/F values across 30° step trials for responsive neurons (n=69 neurons). **(H)** Mean normalized fluorescence traces for one neuron during baseline and response to ipsilateral and contralateral roll steps of varying magnitudes (10°, 20°, 30°). **(I)** Mean peak ΔF/F responses across all responsive neurons for ipsilateral and contralateral rolls of 10°, 20°, and 30° magnitudes. Error bars ± SEM. **(J)** Distribution of directionality indices (Methods) across all responsive neurons.

To image fluorescence, the platform is mounted underneath a water-dipping objective on a two-photon microscope capable of rapid volumetric scanning. All experiments were conducted in complete darkness. A drop of water, held in place by surface tension, sits between the agarose and the objective. Prior to starting an experiment, the platform is adjusted to sit horizontally underneath the objective such that the neurons of interest are in focus. We measured fluorescence across the volume to define a baseline for the neurons. For the step stimuli, we rotated the platform to an eccentric angle (where the neurons of interest are no longer in focus), held the platform at that orientation for 15 seconds, and then returned the platform back to horizontal (Figure 1C). There, we measured the changes in fluorescent intensity in response to our stimulus.

### Vestibulospinal neurons respond reliably to ipsilateral step stimuli

We began by measuring responses to roll tilts at 4 days post-fertilization (dpf) in vestibulospinal neurons (Figure 1E) labelled in transgenic line, *Tg(nefma:GAL4;14xUAS:GCaMP6s)* (Liu et al., 2020). Previous work in larval zebrafish (Liu et al., 2020) and other animals (Fujita et al., 1968) established that vestibulospinal neurons increase their activity as a function of roll tilt angle, with a strong preference for tilts in the direction of the recorded neuron (i.e. a cell in the left hemisphere responds when the left ear is rolled down; henceforth called “ipsilateral roll”). Vestibulospinal neurons therefore provide an excellent test-bed to evaluate new tilt paradigms, such as the step and impulse stimuli we use here.

We mounted the fish parallel to the platform’s axis of rotation to provide both ipsilateral and contralateral roll tilts of 10°, 20°, and 30° (Figure 1C). We detected significant changes in fluorescence relative to each cell’s own baseline (henceforth called “responsive cells”, Methods) in 94% of neurons (67/71 neurons from 10 fish) (Figure 1F, example responsive trace). Importantly, responses were reliable across repeated trials, with a median coefficient of variation across trials of 0.19 (Figure 1F,1G). Consistent with prior reports, responses were direction-dependent. The majority of neurons had a larger response to ipsilateral roll compared to contralateral (Directionality Index = 0.46 ± 0.43) (Figure 1H,1J). Additionally, we found that the strength of responses to roll stimuli increased with the size of the step. For steps in both the ipsilateral and contralateral direction, we observed that the peak response, defined as the mean response within the first second after the end of the stimulus, scaled linearly with the magnitude of the roll step (Figure 1H,1I) (mean slope of responsive neurons= 0.07 ± 0.06 ΔF/F/° ipsilateral, 0.02 ± 0.02 ΔF/F/° contralateral). We conclude that our apparatus can elicit reliable, parametric, and directionally-sensitive responses following roll tilts of different amplitude in vestibulospinal neurons.

### TIPM is robust to extrinsic sources of variability

There are several potential sources of variability that could compromise detection of reliable fluorescent changes following stimulation. First we measured response variation from the following sources: (1) field-of-view movement during imaging and (2) mounting variability. We then measured intrinsic trial-to-trial variability that presumably reflects biological sources such as changes in intraneuronal calcium, spike rate fluctuations, or state of the animal (Schoppik et al., 2008). If the variability observed from trial-to-trial is greater than extrinsic variability, we would conclude that our approach is sufficiently robust.

The dynamic nature of TIPM introduces the potential for the field-of-view of our sample to move during the course of imaging. Sample movement has the potential to cause variability in fluorescence intensity that does not reflect an underlying calcium fluctuation. Qualitatively, we did not observe field-of-view shifts acutely between the baseline recording period and after an eccentric step. We quantified such changes by performing a cross-correlation of each frame before and after the eccentric step. To eliminate signal changes from neuronal fluctuations, we performed this analysis on unprocessed volumes measured in anesthetized fish. The frames before and after the eccentric step were most-highly correlated with each other when they were not shifted relative to each other (mean shift = 0.02 μm in x-axis, -0.01 μm in y-axis; mean correlation without shift = 0.5). Additionally the mean peak fluorescence change after the eccentric step in anesthetized fish was very low (0.08 ± 0.09 ΔF/F anesthetized vs 2.2 ± 1.8 ΔF/F un-anesthetized, n=26 neurons), indicating there is very little variation in the fluorescence signal that results from acute shifts during imaging. We conclude that TIPM as implemented here introduces tolerable levels of sample movement.

Notably, as with any long-term imaging experiment, we did observe that some samples slowly drift in X/Y/Z between trials. We estimate this drift at approximately 1 μm/minute in all axes. These slow shifts can be easily corrected either manually between trials or by *post-hoc* motion correction and so do not introduce appreciable variability into measured responses.

We next addressed the variability due to mounting. For each imaging experiment, larval fish are manually mounted on the galvanometer in agarose in a dorsal-up position. Every attempt is made to minimize roll, pitch, and yaw relative to the axis of rotation, but manual mounting is subject to small variations. These variations would impose linear accelerations that scale with the distance from the center of the axis of rotation. Such shifts would be challenging to quantify and, if large, might compromise longitudinal experiments.

To estimate how much variation in response originated from variation in mounting, we performed a repeated imaging experiment. We mounted and imaged a fish, then removed the fish from agarose and re-mounted the same fish and repeated our imaging protocol (Figure 2A). We were able to reliably identify neurons between the first and second mounts (Figure 2B). To estimate the roll tilt, we calculated the bilateral difference in z-position of the tips of the left and right Mauthner lateral dendrites, and used this to calculate a roll angle of the head. We observed only minor rotation of the baseline position of the fish in the roll axis between the first and second mounts (2.4±1.0°, N=5 fish).

**Figure 2:**
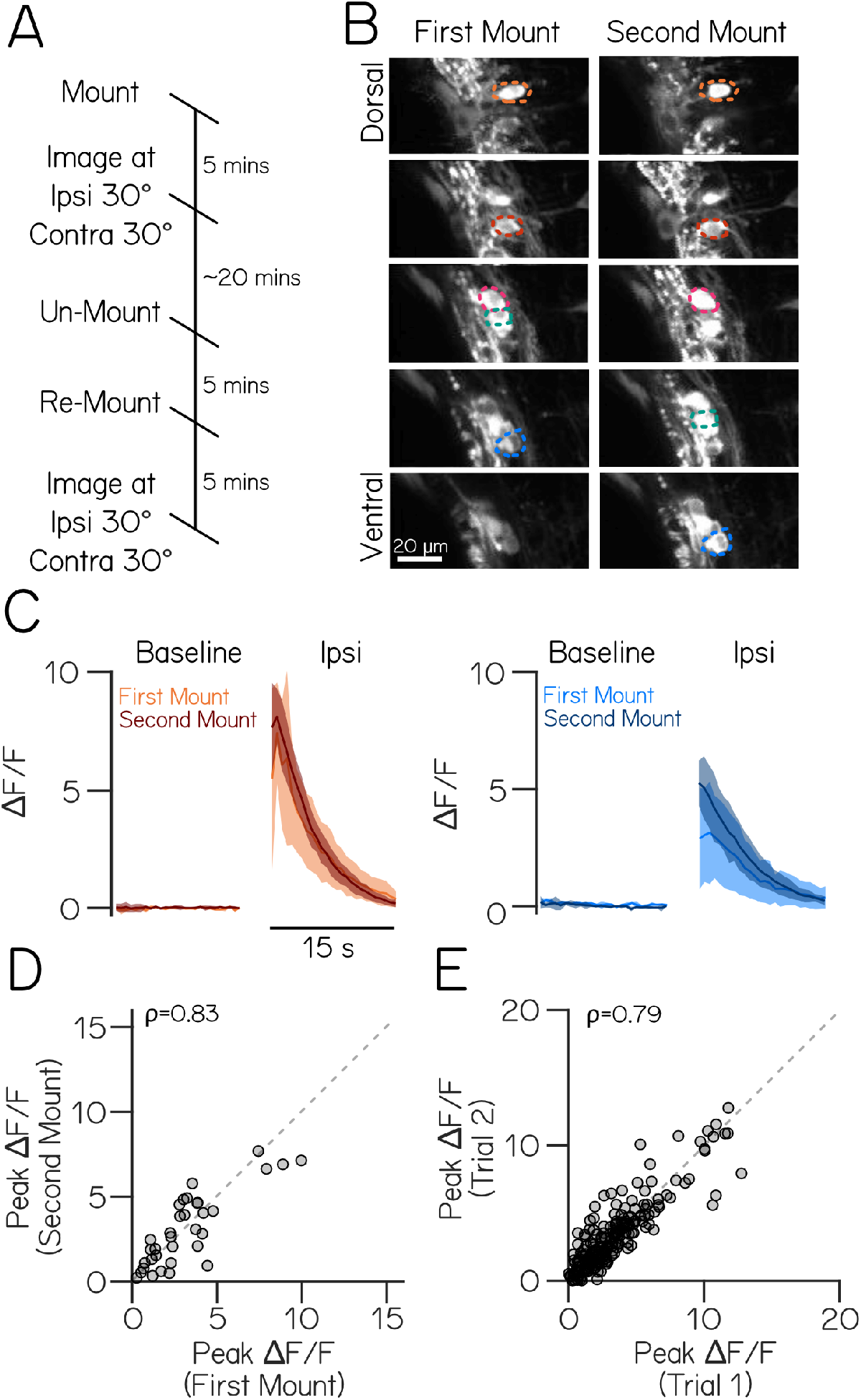
Variability in responses arises predominantly from intrinsic sources. **(A)** Timeline of an experiment of repeated imaging of the same fish across two mounts on the galvanometer. **(B)** Two-photon volumes of vestibulospinal neurons in a *Tg(nefma:GAL4);Tg(UAS:GCaMP6s)* larva during two sequential mounting and imaging experiments. Colored overlays indicate the same neurons located in two separate volumes taken across the two mounts. **(C)** Fluorescence traces from two example cells during the baseline period and following an ipsilateral roll during the first and second mount for two neurons (orange and blue traces correspond to colored neuron overlays). **(D)** Mean peak response during trials in the first mount experiment are strongly correlated with the mean peak response from the second mount (\*ρ*=0.83). **(E)** Peak response on a single experimental trial is strongly correlated with peak response on the subsequent trial within the same experiment (\*ρ*=0.79).

We saw a strong correlation (\*ρ*=0.83) in the response of individual neurons between the first and second mount (Figure 2C-2D). We did not observe a significant shift in responses between the first and second mounts (paired t-test p=0.64, n=34 neurons, N=5 fish), and the responses of the neurons fell nearly along the unity line (slope=0.75 ± 0.18 CI).

To contextualize the magnitude of the mount-to-mount variability we observed, we compared it to trial-to-trial variability. Response correlation between TIPM sessions is comparable to the response correlation between two subsequent trials (Figure 2E) within a single imaging session (\*ρ*=0.79, slope=0.79 ± 0.09 CI, n=34 cells, N=5 fish). These data suggest that most of the variability in response magnitude we see mount-to-mount reflects inherent variability.

Together these experiments establish “best practices” to estimate variability when using TIPM to measure neural responses. We conclude that variability due to our apparatus or mounting are relatively minor concerns for estimating neural response magnitude in our preparation.

### Neural activity imaged after a step reflects the encoding of body tilt prior to the step

Our interpretation of the response rests on the assumption that the activity observed *after* the platform returns to the horizon primarily reflects the activity of the neuron at the eccentric position. To test this assumption, we compared the response of neurons at eccentric angles to that after a step returning the fish to horizontal. In this experiment, we presented the same 30° roll stimulus to fish while measuring activity first upon return to the horizontal plane as previously described (Figure 3A, black), then on subsequent trials measuring activity directly at the eccentric 30° angle (Figure 3A, magenta). Because the light path to the neuron changes as a function of eccentricity, we normalized fluorescence to a baseline stack taken at either the horizontal or the eccentric angle while the fish was anesthetized. We compared fluorescence in vestibulospinal neurons in the first second upon return to horizontal to the responses of the same neurons in the last second of the eccentric step. Neural responses at eccentric angles were closely correlated (\*ρ*=0.94) with the responses measured at horizontal (Figure 3B,3C). The strong similarity in response amplitude supports two conclusions: First, that the response of the neuron upon return to the horizontal is indeed a reasonable proxy for a neuron’s activity at an eccentric angle in the moment just prior to the return step. Second, by inference, larval zebrafish vestibulospinal neurons should have comparatively small responses to phasic stimulation.

**Figure 3:**
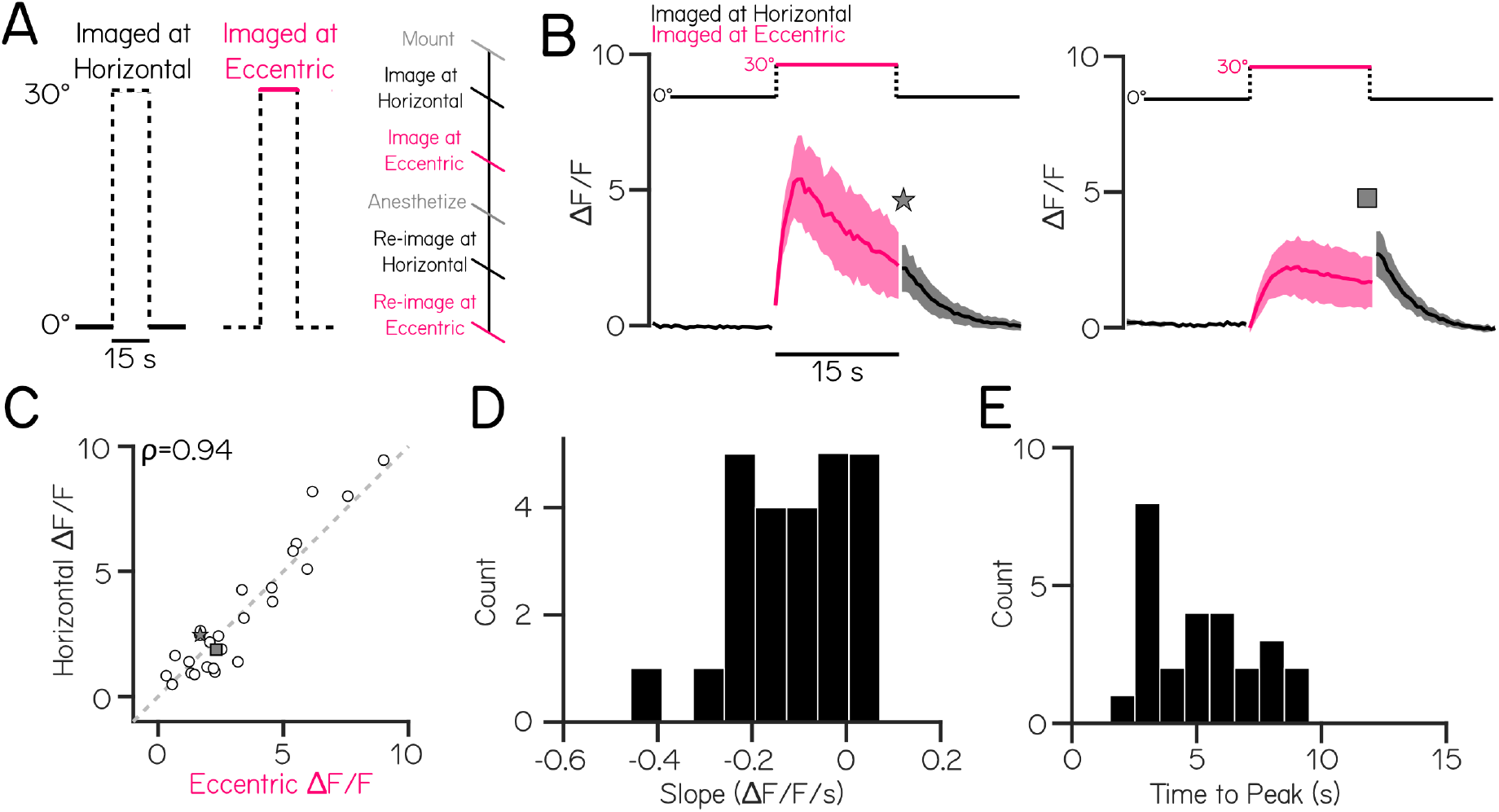
Responses after a step back to horizontal from an eccentric angle are strongly correlated with activity prior to the step. **(A)** Schematic of stimuli used to compare responses after return to horizontal (black) to responses at the eccentric angle (magenta). Solid lines indicate stimulus periods where neurons are in focus, dashed lines indicate stimulus periods where neurons are out of focus. **(B)** Concatenated traces of mean normalized fluorescent responses before, during, and after a ipsilateral roll step of 30° for two example neurons with differing response temporal dynamics (fast-decay on left, slow-decay on right). **(C)** The mean fluorescence during the last second of the eccentric step is strongly correlated with the mean fluorescence during the first second upon return to horizontal for all neurons (n=26 neurons) (\*ρ*=0.94). Example fast and slow-decay neurons in (B) identified with a star and square symbol. **(D)** Distribution of calcium response decay slopes during the eccentric step for all neurons. **(E)** Distribution of times to peak normalized fluorescence during the eccentric step for all neurons.

Notably, while measuring fluorescent intensity at the eccentric angle, we observed quite different dynamics among vestibulospinal neurons. The most striking variation was in the decay-rate of the fluorescent intensity (Figure 3D). Some neurons had a distinct peak followed by a fast-decay (Figure 3B, left), while others had plateau-like responses that had little to no decay over the 15 second hold (Figure 3B, right). In all neurons measured, fluorescent intensity reached its peak within 10 seconds of being at the eccentric angle (median 5.2 s) (Figure 3E). We conclude that for vestibulospinal neurons a 10 second step would be sufficient to ensure accurate detection of the peak response upon return to horizontal (correlation between eccentric calcium response at its peak and at 10 seconds, *ρ*=0.91). As this value will vary between neuronal populations, preliminary experiments to set the optimal window should be done for each new population of interest. By adjusting the length of the TIPM eccentric step to one’s own experimental goals and observed calcium dynamics, the experimenter can use the return to horizontal response as a proxy to measure the magnitude of either the peak or steady-state calcium responses. Taken together, measuring fluorescence upon return to horizontal can be used to accurately extrapolate information about the neuron’s response at the eccentric angle. Further, while imaging at horizontal alone can not provide information about temporal dynamics, we demonstrate here how TIPM can be modified to allow for imaging at an eccentric angle to study variations in response dynamics within a population of neurons.

### Vestibulospinal neurons respond weakly to impulses of angular acceleration

To measure the impulse response of vestibulospinal neurons, we delivered rapid roll steps to 10°, 15°, or 30° and then back to horizontal in *<*13 ms (Figure 4A,4B; Table 1). We observed significant changes in fluorescence to the impulse stimulus in a moderate fraction (35.4%) of neurons (n=22/62 neurons from N=6 fish). The average peak fluorescence observed to the impulse stimulus was small (0.53 ± 0.39 ΔF/F for an ipsilateral 30° stimulus) compared to the response to the tilt stimulus. Impulse responses are more variable across trials than responses to the step stimuli (median coefficient of variation = 0.76 vs 0.19) (Figure 4D). Unlike responses to steps of different amplitudes, peak fluorescent intensity did not vary systematically with the magnitude of the impulse (slope of peak fluorescence = 0.002 ± 0.01 ΔF/F/° ipsilateral, -0.002 ± 0.02 ΔF/F/° contralateral) (Figure 4F). Additionally, impulse responses did not show a consistent directional-preference. Most neurons responded equally strongly to ipsilateral and contralateral steps (Directionality Index = 0.10 ± 0.37) (Figure 4E,4G).

**Figure 4:**
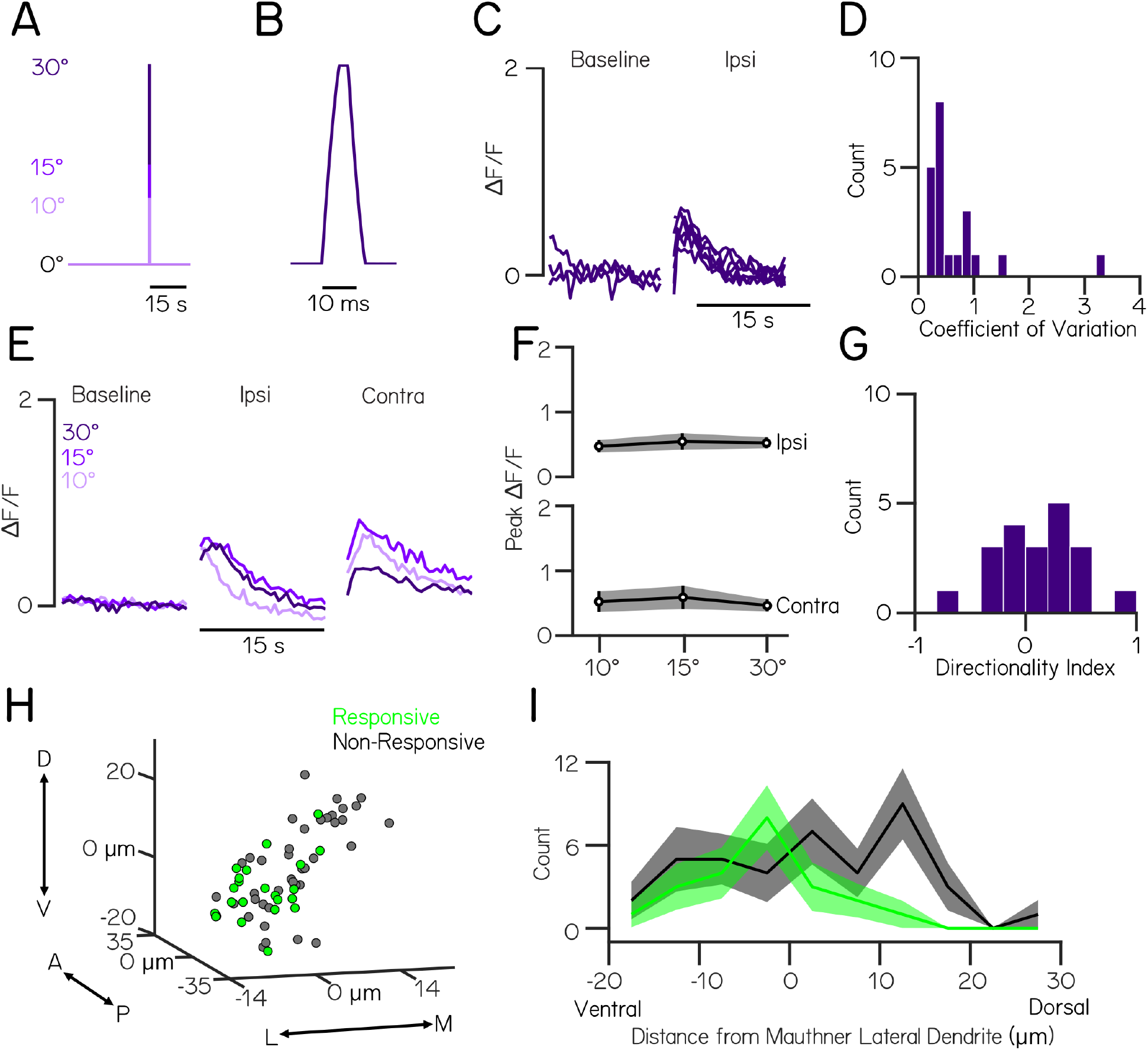
Ventral vestibulospinal neurons respond to impulse stimuli in a non-directional, magnitude-independent manner. **(A)** Voltage-trace corresponding to feedback from galvanometer during a 10°, 15° and 30° impulse step to the left. **(B)** Inset of feedback trace during the impulse step to 30°. **(C)** Normalized fluorescence traces for all trials of one neuron during baseline and response to an ipsilateral 30°impulse step. Note the lower scale on the vertical axis relative to Figures 1-3 **(D)** Distribution of coefficients of variation of peak fluorescent intensity across 30° step trials for responsive neurons (n=22 neurons) **(E)** Mean normalized fluorescence traces for one neuron during baseline and response to an ipsilateral and contralateral impulse step of varying magnitudes (10, 20, 30°). **(F)** Mean peak ΔF/F responses across all responsive neurons for ipsilateral and contralateral impulse steps of 10, 15, and 30° magnitudes. Error bars ± SEM. **(G)** Distribution of directionality indices (Methods) across all responsive neurons. **(H)** Spatial location of impulse-responsive (green) or non-responsive (gray) vestibulospinal somata relative to the Mauthner neuron lateral dendrite in μm. **(I)** Distribution of dorsoventral position of vestibulospinal neuron bodies relative to the Mauthner lateral dendrite in μm for impulse-responsive (green) and non-responsive (gray) neurons.

We asked whether there was topographical organization to these responsive neurons within the lateral vestibular nucleus. While non-responsive neurons are found distributed evenly throughout the lateral vestibular nucleus, neurons with a significant response to the impulse stimulus are located more ventro-laterally (mean dorsoventral position relative to Mauthner lateral dendrite = -3.6 ± 7.0 μm responsive neurons vs 2.2 ± 11.0 μm non-responsive neurons; p=0.03) (Figure 4H,4I). There may therefore be topographic differences in innervation by VIII^th^ nerve afferents that relay phasic vestibular inputs relative to tonic inputs, such that impulse-responsive afferents target only a subset of ventro-lateral vestibulospinal neurons.

Taken together, our data argue that vestibulospinal neurons are more sensitive to the tonic component of the step stimulus than to an impulse stimulus. We infer that the response of vestibulospinal neurons predominantly reflects static encoding of body tilt.

### The utricle is indispensable for the bulk of vestibulospinal neuron responses

Loss-of-function experiments assaying both behavior (Bianco et al., 2012; Ehrlich and Schoppik, 2019; Mo et al., 2010) and neuronal responses (Liu et al., 2020) support the proposal that in larval zebrafish, the bulk of the vestibular response is derived from a single vestibular end-organ: the utricle. However, TIPM is inherently multimodal, and might activate other systems in addition to the utricle. Angular accelerations can be transduced by the semicircular canals. While the semicircular canals are too small to be activated under natural conditions (Beck et al., 2004; Lambert et al., 2008), they can be activated by sufficiently strong stimuli in comparably small vertebrates (Branoner and Straka, 2014). Translational forces along the body might be encoded by the lateral line (Dijkgraaf, 1963). Finally, pressure along the body might be encoded by the trigeminal system (Ribera and Nüsslein-Volhard, 1998). As vestibulospinal neurons are known in other animals to receive a wide variety of multimodal inputs (Sarkisian, 2000) we sought to clarify the role of utricular sensation.

We adopted a genetic loss-of-function approach to assay the contribution of the utricle to vestibulospinal responses. Mutants in *otogelin*, also known as *rock solo* fish (Whitfield et al., 1996), do not form a utricular otolith in the first 10 days (Roberts et al., 2017). *otogelin* is selectively expressed in the inner ear (Stooke-Vaughan et al., 2015), avoiding off-target confounds. We tested the responses of *rock solo* mutants to both a 20° step and impulse stimulus. We provided both ipsilateral and contralateral impulses; as we previously observed no systematic differences we aggregated the data to assay responses.

We observed that vestibulospinal responses to ipsilateral roll steps in *rock solo* fish were severely compromised. The *rock solo* mutants were less likely to show significant changes in fluorescent intensity following a 20° ipsilateral step compared to their wildtype siblings (96% responsive WT, n=24/25 neurons from N=3 fish; 42% responsive mutants, n=13/31 neurons from N=3 fish). When there were supra-threshold responses, the magnitude of peak fluorescence in mutants was strongly attenuated (2.7 ± 2.3 ΔF/F WT vs 0.21 ± 0.24 ΔF/F mutants, 3-way ANOVA Interaction Effect<0.001, post-hoc test p=5.9×10^−8^) (Figure 5A,5B).

**Figure 5:**
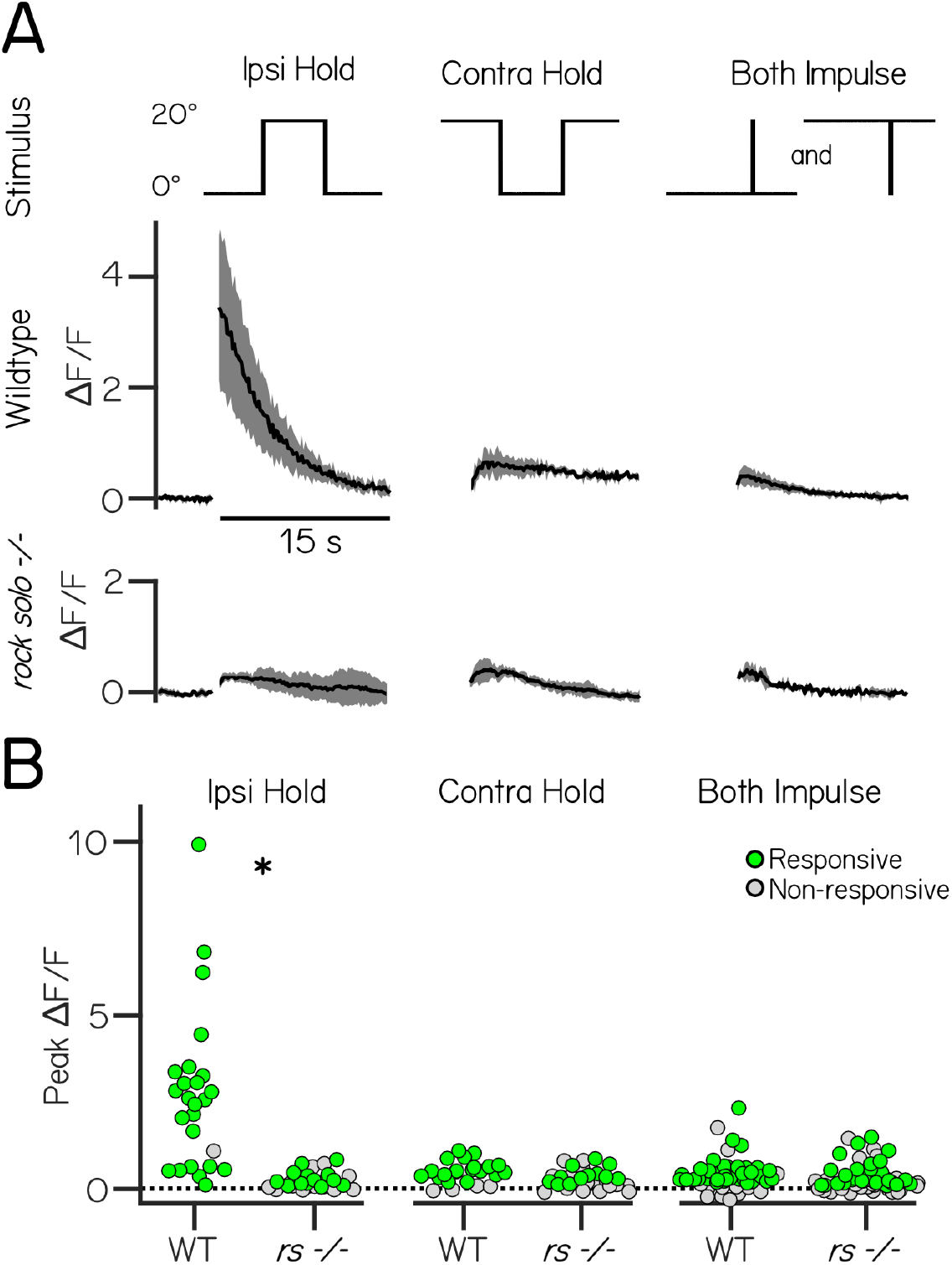
The utricle is indispensable for responses to ipsilateral steps, but not contralateral steps or impulses. **(A)** Example responses (mean ± SD) to 20° step and impulse stimuli (top row) from vestibulospinal neurons in wildtype and *rock solo* mutants. **(B)** Peak fluorescence responses to ipsilateral step, contralateral step, and impulse steps in both directions in wildtype and *rock solo* mutants in responsive (green) and non-responsive (gray) neurons.

In contrast, responses were not significantly different between wildtype siblings and *rock solo* mutants following contralateral steps (0.38 ± 0.41 ΔF/F WT vs 0.26 ± 0.30 ΔF/F mutant), nor were responses significantly different to impulse steps (0.42 ± 0.48 ΔF/F WT vs 0.32 ± 0.39 ΔF/F mutant; post-hoc test p=0.99) (Figure 5A,5B) We conclude that contralateral eccentric and impulse responses are predominantly driven by extra-utricular sources. Following both contralateral steps and impulse stimuli, we observed a decrease in the fraction of neurons that responded to the stimulus in *rock solo* fish (Contralateral step = 72% responsive WT vs 39% responsive mutant, Impulse = 72% responsive WT vs 44% responsive mutant). Changes to the fraction of neurons that have supra-threshold responses reflect an increase in variability of baseline calcium fluctuations in *rock solo* mutants (baseline SEM: 0.019 WT vs 0.032 rock solo), consistent with electrophysiological observations (Hamling et al., 2021; Liu et al., 2020).

We conclude that the changes in fluorescence we observe in vestibulospinal neurons following ipsilateral body tilts predominantly reflects utricular transduction.

### Vestibulospinal neuron responses develop systematically

A distinct advantage of TIPM is its minimally-invasive nature. As such, it is well-suited for experiments that require monitoring the same neurons across multiple timepoints. We asked if TIPM could detect developmental changes in individual vestibulospinal neurons on two different days. Prior behavioral work established that larval zebrafish use vestibular information to balance and locomote in different ways at 4 and 7 dpf (Ehrlich and Schoppik, 2017, 2019). We therefore picked 4 and 7 days to assay for differences in body tilt-evoked responses.

We imaged fluorescence after return to horizontal from 10°, 20°, and 30° step stimuli in the same fish at two ages: 4 and 7 dpf. We were able to reliably identify the same neurons across imaging sessions (Figure 6A). Peak calcium responses within the same neuron were correlated (\*ρ*=0.51) between 4 and 7 dpf (n=71 cells, N=10 fish)(Figure 6B), but were more variable than neurons in repeated imaging sessions performed on the same day (Figure 6B, gray bar) suggesting developmental changes in neuronal encoding. We observed that responses were more variable in our second imaging session (7 dpf, median CV=0.29) than the first (4 dpf, median CV=0.19); we therefore chose to compare the slope of peak fluorescence responses (“Roll Sensitivity,” a common metric of sensory encoding capacity (Lannou et al., 1979)) of neurons across development, instead of a metric like mutual information that takes response variability into account (Quiroga and Panzeri, 2009). Across all neurons that responded to the stimulus at either age, calcium response sensitivity to ipsilateral eccentric rolls strengthened between 4 and 7 dpf (mean slope = 0.07 vs 0.10 ΔF/F/°, Repeated Measures ANOVA post-hoc test p=0.002; n=70 cells), and sensitivity to contralateral roll did not decrease significantly (mean slope = 0.02 vs 0.01 ΔF/F/°, p=0.09). Our data suggest that the population of vestibulospinal neurons improves its ability to encode eccentric roll tilts during this developmental window.

**Figure 6:**
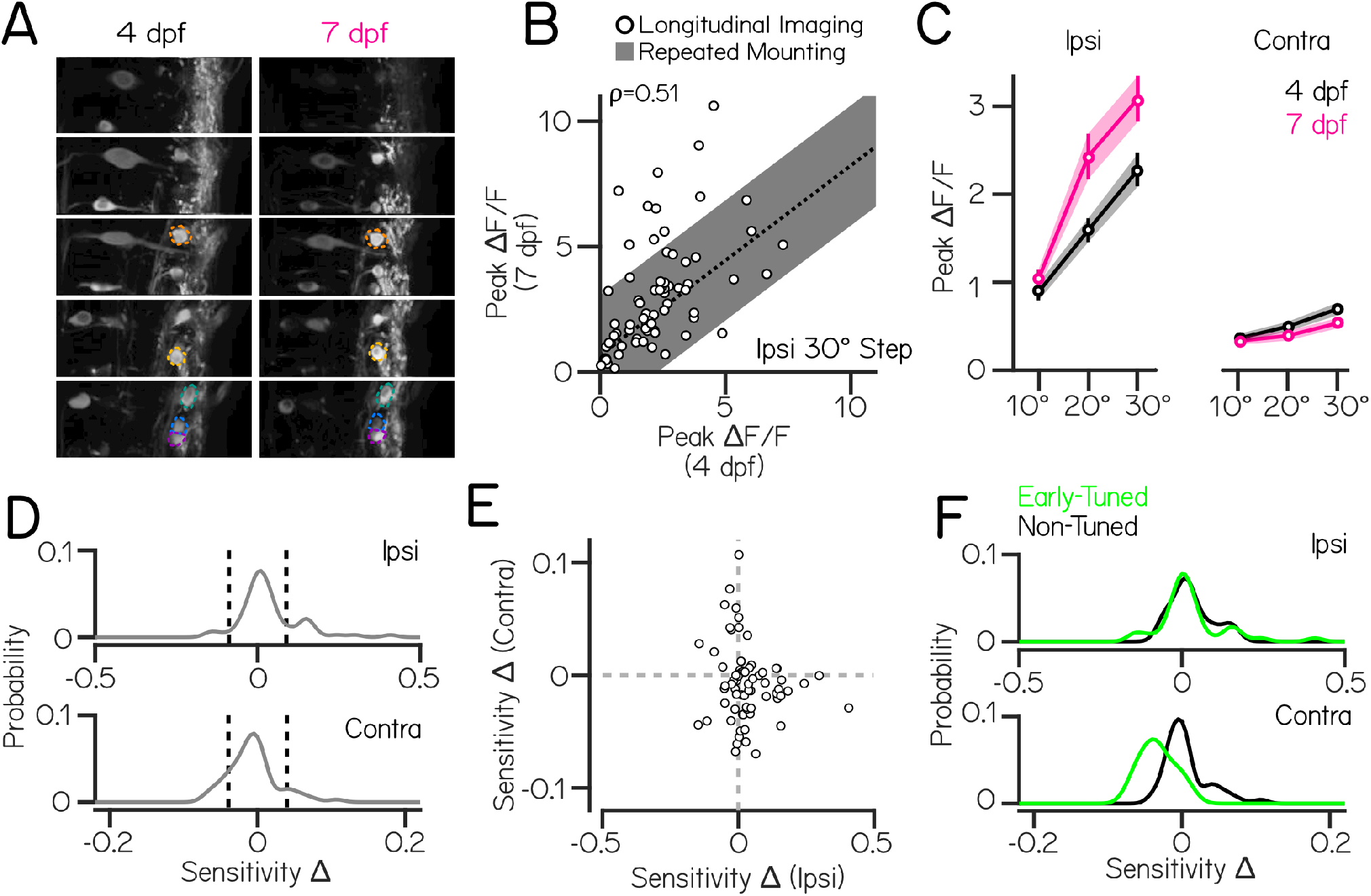
Longitudinal imaging suggests systematic changes in the complement of pre-synaptic inputs in developing vestibulospinal neurons. **(A)** Two-photon volumes of vestibulospinal neurons in a *Tg(nefma:GAL4);Tg(UAS:GCaMP6s)* larva during longitudinal imaging experiments at 4 and 7 dpf. Colored overlays indicate the same neurons located in the volume across the two timepoints. **(B)** Mean peak calcium response of all neurons at 4 dpf are correlated with their mean peak response at 7 dpf (\*ρ*=0.51), but is more variable than expected by remounting alone (fit line ± S.D. of residuals from repeated mounting experiment in Figure 2D). **(C)** Mean peak ΔF/F responses across all responsive neurons for ipsilateral and contralateral rolls of 10, 20, and 30° magnitudes at 4 (black) and 7 dpf (magenta). Error bars ± SEM. **(D)** Probability distributions of longitudinal changes in ipsilateral (top) and contralateral (bottom) calcium response sensitivity (Methods) (n=70 cells). Dashed vertical lines represent cut-offs for significant sensitivity change. **(E)** Longitudinal sensitivity changes in individual neurons across ipsilateral and contralateral roll tilts. **(F)** Longitudinal sensitivity changes for ipsilateral (top) and contralateral (bottom) roll tilts split by neurons that have significant magnitude-dependent responses to roll steps in that direction at 4 dpf (“Early-Tuned,” green) versus non-tuned at 4 dpf (“Non-Tuned,” black).

Additionally, beyond effects on the population, our longitudinal imaging paradigm allowed us to ask 1) how an individual neuron’s sensitivity to ipsilateral or contralateral eccentric roll angle changed across development and 2) whether these changes are systematically patterned. First, we investigated whether developmental changes within the population of vestibulospinal cells was homogeneous. To do so, we examined the distribution of developmental sensitivity changes in individual cells. We then identified cells with a significant change in roll sensitivity by comparing the observed change in a cell’s sensitivity to a cutoff generated from sensitivity changes in age-shuffled data (Methods). We found that individual cells had heterogeneous and asymmetric sensitivity changes to ipsilateral rolls (Figure 6D, top). In response to ipsilateral rolls, individual vestibulospinal cells experienced either no change in sensitivity (53/70 cells) or a significant increase in sensitivity (14/70 cells) but very rarely experienced a significant decrease in ipsilateral sensitivity between 4 and 7 dpf (3/70 cells). This asymmetry explains the overall increase in ipsilateral responses observed across the whole population at 7 dpf. In comparison, the distribution of contralateral sensitivity changes was heterogeneous and approximately symmetric (Figure 6D, bottom), with most cells experiencing no significant change (51/70 cells) and comparable numbers experiencing a significant sensitivity increase (8/70 cells) or decrease (11/70 cells). Together, these findings allow us to conclude that the vestibulospinal population is not homogeneous in how tilt responses develop. Specifically, a majority of cells do not change between 4 and 7 days while small sub-populations either increase or decrease their sensitivity in a directional-dependent manner.

We then asked whether the developmental changes an individual cell experiences in the ipsilateral and contralateral directions are correlated; we found that there was no significant correlation between sensitivity changes across these two directions (Figure 6E) (\*ρ*=-0.14, p=0.25). Together with previous findings, these data support the existence of different patterns of functional development occurring within the vestibulospinal nucleus that are not coordinated between response directions. While ipsilateral responses strengthen and rarely weaken, the same pattern does not apply to contralateral responses. Additionally, developmental change in one direction does not predict change in the opposing direction. The lack of correlation observed between ipsilateral and contralateral developmental changes suggests that the refinement of these two directions are driven by separate mechanisms.

To explore potential mechanisms for systematic changes in response properties across development, we attempted to predict how ipsilateral and contralateral roll sensitivity within a neuron would change based on the responses we observed at 4 dpf. We asked whether neurons with a significant, magnitude-dependent response to roll stimuli early in development (“Early-Tuned”) would selectively strengthen or weaken their responses as they develop. For ipsilateral stimuli, there was no significant difference between sensitivity change distributions when split between early-tuned (42/70 neurons) and non-tuned cells (Figure 6F, top) (two-sample Kolmogorov-Smirnov test, p=0.99). In contrast, we found that neurons that were early-tuned for contralateral stimuli (24/70 cells) had a significantly different distribution of developmental sensitivity changes compared to non-tuned neurons (Figure 6F, bottom)(two-sample Kolmogorov-Smirnov test, p=2.06×10^−7^). Specifically, early-tuned contralateral neurons were only observed to decrease (11/24 neurons) or have no change (13/24 neurons) to their contralateral sensitivity between 4 and 7 dpf, and never strengthened their contralateral responses. Neurons that did not have significant contralateral responses at 4 dpf either strengthened their contralateral sensitivity between 4 and 7 dpf (8/46 neurons) or experienced no change (38/46 neurons). We can therefore conclude that contralateral, but not ipsilateral, sensitivity changes are patterned by a neuron’s responses early in development. Such inferences are new to the vestibular field and are made possible by our ability to follow the same neurons over multiple days. We propose that our novel approach is therefore well-suited for discovering biologically-relevant changes responsible for improvements to neuronal control of posture as animals develop.

## Discussion

Here we report a new method, Tilt In Place Microscopy, to measure neuronal responses following vestibular stimulation. To test TIPM, we mounted a larval zebrafish on a rotating platform (mirror galvanometer) and measured fluorescence as the fish returned from an eccentric tilt. Consistent with prior work (Peterson, 1970; Rovainen, 1979), we observed reliable responses that vary with tilt magnitude. We tested the reproducibility of our method by mounting the same fish repeatedly, finding that TIPM produced little change in the strength and reliability of neuronal responses. By imaging the same neurons both at an eccentric angle and at the horizon, we confirmed that the response reflected the steady-state activity while tilted. We next delivered impulse steps, and discovered topographically-organized responses. Consistent with other work, vestibulospinal neurons in mutant zebrafish without utricles responded minimally following tilts. Finally, we measured activity from a set of vestibular neurons at both 4 and 7 days post-fertilization and reveal systematic changes in sensitivity and selectivity across time. Below we compare TIPM to other approaches/apparatus, discuss limitations and potential extensions, and contextualize our findings.

### Comparison to other apparatus/approaches

Other groups have tackled the challenge of imaging neural activity while measuring vestibular responses; each approach requires considerable technical expertise and financial resources. One approach involved adapting a powerful technique, the optical trap, to directly displace the utricle (Favre-Bulle et al., 2018). Another approach cleverly miniaturized the hardware so as to permit an entire light sheet microscope to rotate stably (Migault et al., 2018). Together, these apparatus allowed these investigators to characterize vestibular responses across an entire vertebrate brain – a considerable advance. Recently, another group developed a rotating stage that allowed them to examine the vestibular periphery (Tanimoto et al., 2022). Finally, a complementary approach used for *in vivo* electrophysiology may be compatible with imaging provided that the microscope was sufficiently small: mounting the preparation on an air-bearing sled (Liu et al., 2020). All four solutions require a familiarity with optical physics and/or engineering to implement and calibrate. All four require specialized and expensive hardware.

In contrast, TIPM offers a number of advantages. It is comparatively low-cost and compatible with any microscope (multiphoton or otherwise) with a long working distance objective that can accommodate a simple rotating platform. Importantly, the only requirement to control the stimulus is the ability to deliver an analog voltage to control the galvanometer and a digital pulse for calibration. We propose that TIPM’s simplicity, flexibility, and low cost facilitates the study of the neuronal encoding of vestibular responses.

### Limitations of TIPM

The design choices we made facilitate certain experiments, but are not without their own trade-offs. TIPM was designed to image the response to tilts at one particular orientation, to facilitate comparison to baseline measurements so crucial for imaging fluorescence from calcium indicators. Responses must be therefore measured *after* stimulation is complete. Further, we chose to move the preparation relative to a stationary microscope objective. The light path to the same neuron will change with each change in orientation. Consequentially, comparisons across orientations (as in Figure 3) requires normalization to a baseline that accounts for different scattering such as an anesthetized baseline (Methods). The inability to measure during stimulation and the challenge of comparing across orientations means that our method is likely incompatible with a sinusoidal rotation.

The vestibular periphery has been been modeled as an linear, time-invariant system (Laurens et al., 2017). By and large, the measurements underlying this powerful framework are derived from sinusoidal stimulation at different frequencies while recording neuronal responses across vestibular areas. However, sinusoidal stimulation is not the only way to measure the response of a linear, time-invariant system. Impulse stimuli (e.g. Figure 4A) contain power at a wide range of frequencies. Such “click” stimuli are common in characterizing auditory responses and the head impulse test is common place during clinical evaluation of semicircular canal function (Halmagyi et al., 2017). Similarly, step stimuli such as we have used here allow evaluation of the DC component (i.e. the steady-state response to gravity at a particular orientation). While TIPM as presented here is incompatible with sinusoidal rotation, we propose that evaluating the responses to impulses (Figure 4) and steps (Figure 1) as we have here will serve comparably for linear systems analysis of the vestibular system.

Unlike other systems that can rotate a full circle, TIPM is constrained to a smaller range of angles due to three factors. First, the galvanometer itself can only rotate 40°. Second, our preparation relies on surface tension to keep the water between the sample and the objective. In practice rotations greater than 40° relative to the horizon risk spilling the water. Finally, steric considerations limit the achievable rotation. To rotate 90°, the platform would have to be sufficiently narrow so as to fit entirely within the working distance of the objective (2 mm) Such a narrow platform would be unwieldy to mount and hold too little water. We therefore do not believe our apparatus will be able to tilt much beyond what we report here. Experiments that require a wider range of angles are better performed on apparatus that can rotate more.

### Ways to extend TIPM

For imaging experiments, the choice of indicator and field of view set fundamental limitations in time and space. Here we used a slow calcium indicator (GCaMP6s) to measure neuronal activity. All our estimates of vestibular response are convolved with the spike-to-calcium kernel (Chen et al., 2013). This low-pass filter constrains our ability to measure vestibular responses regardless of whether stimuli are sinusoidal, impulses, or steps. Recent advances in genetically-encoded voltage indicators suggest that fluorescent imaging of membrane potential is on the horizon (Böhm et al., 2022), or perhaps here (Liu et al., 2022). As TIPM delivers rapid changes to tilt, and is straightforward to integrate with advanced microscopes, we anticipate that it will be ideal for voltage imaging experiments. Similarly, TIPM translates readily to microscopes with wider fields of view facilitating “whole-brain” approaches.

TIPM can accommodate a wide variety of existing hardware to accommodate *in vivo* imaging in different preparations. While we used a mirror galvanometer, any device that can rapidly and precisely rotate away from and back to a given angle will work. Small direct-drive rotation mounts such as Thorlabs DDR25, or larger options such as Newport’s RGV100 series offer rapid and precise rotation and allow for larger loads than the mirror galvanometer. A low-cost option is similarly available by substituting a DC stepper motor and a driver with micro-stepping capability to permit smooth acceleration. Both options would also permit compatibility with current platforms for *in vivo* imaging in *Drosophila* (Aragon et al., 2022) and *Caenorhabditis* (Smith et al., 2022). Naturally, our approach is compatible with head-mounted microscopes (Aharoni and Hoogland, 2019; Zong et al., 2022) and stably-mounted high-density probes of electrical activity (Steinmetz et al., 2021) in rodents. We anticipate that labs looking to adopt TIPM will select the device that best rotates their existing preparation.

TIPM can be easily extended to permit measuring tail, fin, and/or eye movements in zebrafish. Similar adaptations allow for measurement of leg/wing movements in other animals. First, it is necessary to replace the mirror galvanometer with a transparent platform. A camera mounted below the apparatus can measure the tail bends and eye movements in the horizontal plane with freely available software such as Stytra (Štih et al., 2019). To measure eye movements in the torsional plane, a glass coverslip can be glued perpendicular to the plane of the slide. A camera can then measure torsional eye movements that follow pitch tilts, as done in (Bianco et al., 2012). As tilt stimuli reliably elicit compensatory postural and ocular behaviors, such apparatus would provide valuable context to the measures of neuronal activity we report here.

### Insights into encoding of roll tilt by developing vestibulospinal neurons

The responses to roll tilts in vestibulospinal neurons reported here largely agree with and extend prior reports, bolstering confidence in TIPM, and go on to describe novel development findings that have not previously been observed in vestibular nuclei in any species. We see much stronger responses to ipsilateral than to contralateral steps, replicating findings from larval zebrafish (Hamling et al., 2021; Liu et al., 2020) and other animals (Peterson, 1970; Rovainen, 1979). Mature vestibulospinal neurons are thought to integrate static otolithic information and dynamic information from the semicircular canals (Sarkisian, 2000). We observe minimal impulse responses, consistent with prior reports that larval zebrafish semicircular canals are too small to transduce angular accelerations (Beck et al., 2004; Lambert et al., 2008), and with our observation that loss of the utricle profoundly decreases the responses to tilts. The small impulses we do see are consistent with more recent work in *Xenopus* that provided considerably larger accelerations to reveal canal-mediated responses (Branoner and Straka, 2014). We conclude that the measurements of neuronal activity in vestibulospinal neurons that we performed here to test our apparatus are likely a reasonable measure of tilt sensitivity.

We report systematic changes in neuronal responses from the same vestibulospinal neurons measured at two different ages. Our choice of age was guided by prior reports showing behavioral differences in vestibular-mediated locomotion developing between days 4 and 7 post-fertilization (Ehrlich and Schoppik, 2017, 2019). While two timepoints are too few to truly define a developmental trajectory, we observed that across the population, ipsilateral responses strengthened. Such a trend is consistent with the mature tuning for ipsilateral roll reported for vestibulospinal neurons (Peterson, 1970; Rovainen, 1979). By following the same cells over time, TIPM longitudinal imaging also allows us to make new observations about patterns of functional vestibular development. We found that response sensitivity changes do not appear random but instead follow structured patterns which, for some response types, are related to response properties early in development. Our analyses point the way forward to use early response properties to predict how neurons will change across development.

### Conclusion

From birth to death, an animal’s sense of gravity and other accelerations profoundly shapes its physiology (Yates et al., 2013) and journey through the world (Angelaki and Laurens, 2020). Perhaps unsurprisingly, studies of the vestibular system have informed nearly every aspect of modern systems-level neuroscience (Goldberg et al., 2012a). Advances in imaging neuronal activity have similarly shaped modern neuroscience. Others have brought imaging and vestibular stimulation together with custom microscopes (Favre-Bulle et al., 2018; Migault et al., 2018; Tanimoto et al., 2022), but adoption requires considerable expertise and financial resources. Here we describe and validate a novel apparatus/analysis approach we call TIPM to image neuronal responses to body tilts. TIPM is comparatively easy to implement, compatible with a large set of existing microscope designs, low-cost, non-invasive, extensible to a wide variety of preparations, and compatible with longitudinal measurements. We support this claim by confirming and extending our understanding of tilt representation by developing vestibulospinal neurons in the larval zebrafish. Specifically, we observed preferential sensitivity to tonic stimulation, rough topographic organization, tractable levels of extrinsic variability, and systematic changes across early development. While not without trade-offs, we hope that the simplicity and broad compatibility of TIPM will democratize the study of the brain’s response to destabilization, particularly across development.

## Acknowledgments

Research was supported by the National Institute on Deafness and Communication Disorders of the National Institutes of Health under award numbers DC017489 and DC019554, the National Institute of Neurological Disorders and Stroke under award number NS125280, by the Leon Levy Foundation, by the Irma T. Hirschl/Monique Weill-Caulier Trust, by the Brain Research Foundation, and by the Dana Foundation. On Elizabeth M. C. Hillman’s suggestion, Marie R. Greaney was the first person (to our knowledge) to mount a larval zebrafish on a mirror galvanometer. She and David E. Ehrlich performed pilot experiments used to develop the paradigm described here. The authors would like to thank Jong-Hoon Nam, Martha Bagnall, and the members of the Schoppik and Nagel labs for their valuable feedback and discussions.

## Author Contributions

Conceptualization: KH and DS, Methodology: KH, YZ, FA and DS, Investigation: KH, Visualization: KH Writing: KH and DS Editing: DS, Funding Acquisition: KH and DS, Supervision: DS.

## Author Competing Interests

The authors declare no competing interests.

## Notes

### Competing Interest Statement

The authors have declared no competing interest.

### Summary of Updates

1. Significance Statement added 2. Additional detail added to Methods 3. Figure 6 (Development of vestibulospinal responses) overhauled. All changes made in response to comments from two anonymous reviewers.

